# Off-the-Shelf Engineered Liver Tissue Reverses Acute Liver Failure Without Immunosuppression

**DOI:** 10.1101/2025.10.08.680796

**Authors:** Claudia Raggi, Pascal Lapierre, Marie-Agnès M’Callum, Quang Toan Pham, Silvia Selleri, Radu Alexandru Paun, Nissan Baratang, Chenicka Lyn Mangahas, Basma Benabdallah, Gaël Moquin-Beaudry, Jiwoon Park, Dorothée Dal Soglio, Yves Théoret, Robert E. Schwartz, Christian Beauséjour, Elie Haddad, Massimiliano Paganelli

## Abstract

There is an urgent need for effective solutions to replace liver functions in patients with acute liver failure (ALF). We describe here a human engineered liver tissue composed of induced pluripotent stem cell (iPSC)-derived liver organoids encapsulated within a non-degradable biomaterial. Unlike most stem cell–derived products, this encapsulated liver tissue (ELT) achieves functional maturation during manufacturing. When transiently implanted into the peritoneal cavity of immunocompetent mice with ALF, the human ELT improves survival, treats hepatic encephalopathy and promotes liver regeneration, without requiring immunosuppression. Based on robust processes and designed to overcome challenges such as foreign body reaction, loss of function and cryopreservation, the ELT does not require vascularization and provides immediate and long-lasting functional replacement. Once the liver regenerated, the ELT is explanted, leaving the subjects cured. Macroencapsulation prevents rejection by shielding the organoids from the host immune system, and minimizes the risk of tumorigenicity. The data shown demonstrate the ELT’s potential to be developed into a safe and effective off-the-shelf treatment that, if validated in upcoming clinical trials, could replace liver transplantation for many patients with ALF.

## INTRODUCTION

Acute liver failure (ALF) is a severe, rapidly progressing syndrome caused by sudden and extensive liver injury that overwhelms the liver’s regenerative capacity. The prognosis is poor, with fewer than half of patients surviving with their native liver.^1^ Liver transplantation is currently the only effective treatment, but it is associated with less favorable outcomes than for other indications,^2,3^ and must be performed within days to prevent irreversible complications.^1,3^ Temporary replacement of liver functions while the patient’s own liver regenerates might eliminate the need for transplant (and associated lifelong immunosuppression) in most cases, changing the fate of patients with ALF.^4^ For more than 20 years, transplantation of primary human hepatocytes (PHH) from healthy donors was anecdotally reported to induce transient improvement in patients with liver failure or metabolic disorders.^5^ However, donor organ shortage, high inter-donor variability, unpredictable efficacy, the requirement for repeated administrations and intensive immunosuppression, have limited the implementation of this approach. Stem cells have emerged as promising candidates to help address some of these limitations. We and others have explored the differentiation of stem cells from various sources into hepatocyte-like cells (HLC).^6,7^ Despite encouraging progress, stem cell–derived HLCs typically remain immature *in vitro*, acquiring full adult hepatocyte functions only after *in vivo* maturation.^8,9^ The recreation of cell-to-cell interactions between HLC and endothelial and mesenchymal cells in liver organoids was shown to induce significant maturation of liver cells.^10,11^ However, such stem cell-derived organoids are still not mature enough to replace liver functions immediately upon transplantation, which makes them unsuitable to treat ALF.^11–13^

Thus, there is an urgent need for effective liver support systems able to replace liver functions in patients with ALF. We developed a mature human liver tissue composed of induced pluripotent stem cell (iPSC)-derived complex liver organoids encapsulated within a non-degradable biomaterial, which can treat ALF in an immunocompetent preclinical model without the need of immunosuppression. We describe here that, unlike most stem cell-derived liver products, such an Encapsulated Liver Tissue (ELT) achieves full functional maturation *in vitro*. When transiently implanted into the peritoneal cavity of mice with ALF, it improves survival, treats hepatic encephalopathy (HE) and accelerates liver regeneration. Macroencapsulation shields the liver tissue from the host immune system, eliminating the risk of rejection, and prevents the leak of potentially residual undifferentiated cells. These unique features make the ELT a promising allogeneic cell therapy product that, pending clinical confirmation, might potentially replace liver transplantation for most patients with ALF.

## RESULTS

### iPSC-derived liver organoids

The ELT is composed of liver organoids, each constituted of hepatoblasts aggregated with mesenchymal and endothelial progenitor cells, all derived from a single human iPSC population (Fig. 1a). Hepatoblasts were generated using our previously described differentiation protocol, which delivers, with high yield (Fig. 1b), a homogeneous population of liver progenitor cells (Fig. 1c,d) capable of differentiating into hepatocytes and cholangiocytes.^7^ Mesenchymal progenitor cells (MPC) fulfilled the defining criteria of mesenchymal stromal cells in terms of morphology, surface markers and multipotency (Supp. Fig. 1a-f).^14^ Endothelial progenitor cells (EPC) showed features suggestive of endothelial colony forming cells (Supp. Fig. 1g-l).^15^ Seeding a defined *ratio* of hepatoblasts, MPC and EPC in microcavity culture plates recreated favorable conditions for self-aggregation and formation of controlled-size liver organoids (Fig. 1e). These organoids rapidly expressed liver-specific markers (Fig. 1f) and performed liver functions, although less efficiently than PHH or even iPSC-derived HLC (Fig. 1g,h). The organoids’ size was strictly controlled (160.7±61.5 µm in diameter; Fig. 1i) to maintain optimal perfusion. In order to ease future clinical translation, all protocols were adapted to be xeno-free and serum-free, and only compounds that have a good manufacturing practices (GMP)-compliant equivalent were used.

**Figure 1.**
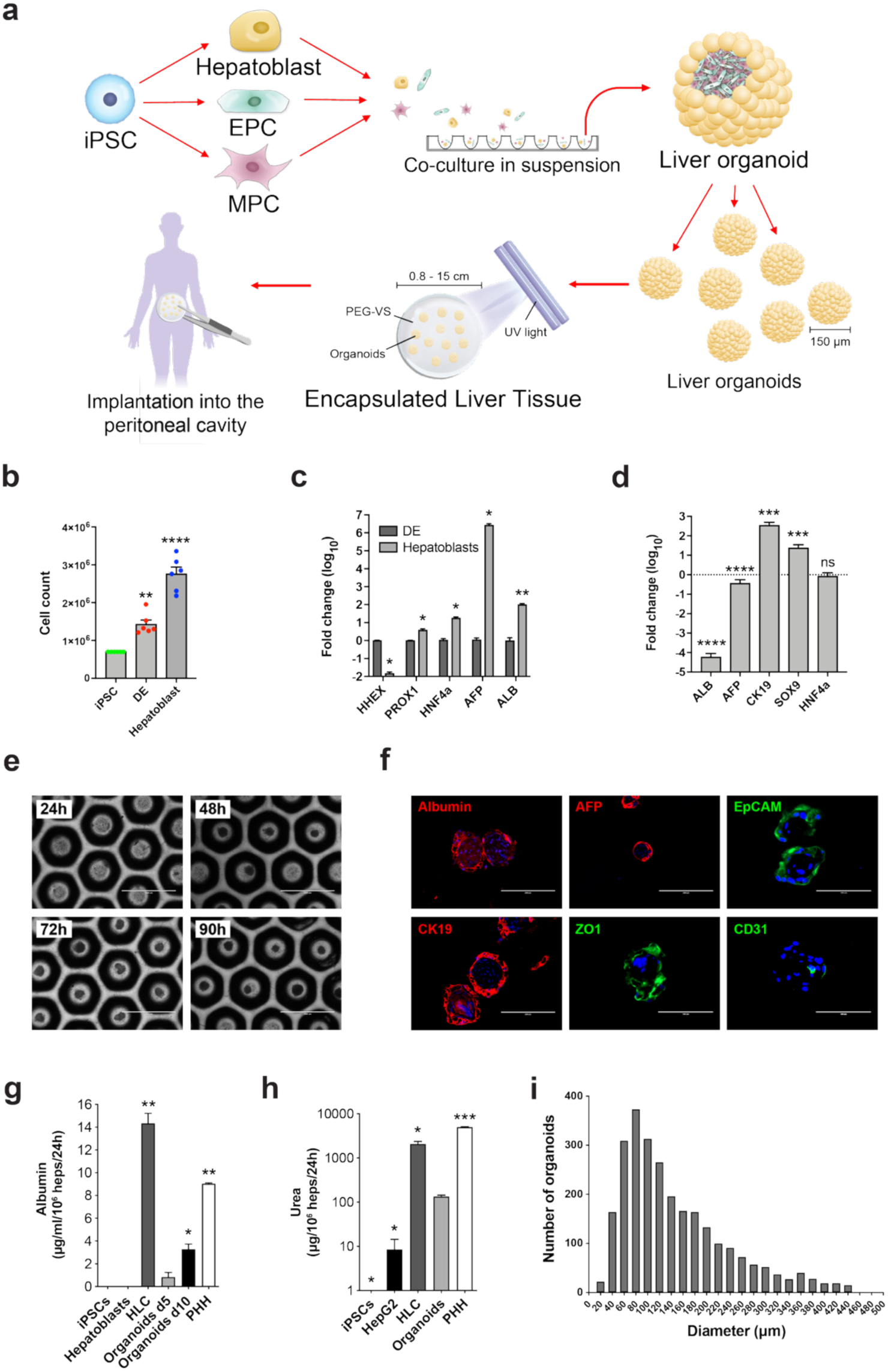
Generation of iPSC-derived liver organoids. **(a)** Schematic of liver organoids organoids and ELT generation. **(b)** The differentiation protocol used allow for increasing yields upon differentiation of iPSC into definitive endoderm (DE) and hepatoblasts (mean ± SE, 6 different batches, paired *t* test, ** *p*<0.01, **** *p*<0.0001 vs. seeded iPSC). **(c)** Liver-specific genes are upregulated upon differentiation from DE to hepatic progenitor cells, with negligible variability between batches (n=4 DE, n=8 hepatoblasts; mean ± SE, Mann-Whitney U test, * *p*<0.05, ** *p*<0.01 vs. DE). **(d)** Compared to the fetal human liver, iPSC-derived hepatoblasts are less mature, with similar expression of *AFP* and *HNF4α*, but lower *ALB* and higher *CK19* and *SOX9* (n=4 for primary fetal hepatocytes, n=8 for hepatoblasts; mean ± SD, one sample *t* test, *** *p*<0.001, **** *p*<0.0001 vs. primary fetal hepatocytes). **(e)** Appearance of liver organoids aggregating within the microcavity plates over 90h after seeding of iPSC-derived hepatoblasts, MPC and EPC (representative images, scale bar 1000 µm). **(f)** Liver organoids rapidly express immature liver-specific markers (albumin, alpha-fetoprotein/AFP, cytokeratin 19/CK19; scale bar 200 µm), adhesion proteins (EpCAM, zonulin-1/ZO1; scale bar 100 µm) and endothelial markers (CD31; scale bar 100 µm; DAPI nuclear staining, representative images of 6 biological replicates). **(g,h)** After only 90h, liver organoids are already capable of performing liver-specific functions, although at levels that are lower than PHH and iPSC-derived hepatocyte-like cells (HLC): **g)** albumin secretion (n=6 biological replicates for organoids day 5, n=3 organoids day 10, n=4 PHH, n=4 HLC; Mann-Whitney test: organoids d5 vs PHH, organoids d10 and HLC, * *p*<0.05, ** *p*<0.01); **h)** urea synthesis (n=6 organoids d5, n=12 PHH, n=3 HLC, HepG2 cells and iPSC; Mann-Whitney test, organoids d5 vs PHH, HLC, iPSC and HepG2, * *p*<0.05, *** *p*<0.001). **(i)** The size of liver organoids is controlled to allow for optimal perfusion of the cells (size distribution, average diameter/organoid, n=2662 organoids from 6 batches/biological replicates).

### Encapsulated Liver Tissue

We optimized a polyethylene glycol (PEG)-based, non-degradable hydrogel to support the survival and maturation of the liver organoids. Multiple organoids were macroencapsulated, as a tissue, within a double layer of 4%, 4-arm, 20 kDa PEG–vinyl sulfone (PEG-VS) hydrogel, which was then photopolymerized under minimal UVA exposure. The resulting 2-mm-thick semi-solid product could be manipulated easily with standard surgical tools while maintaining optimal perfusion (Fig. 2a). Once crosslinked, the non-degradable biomaterial exhibited a Young’s modulus of ∼1.11 kPa, comparable to decellularized human liver (1.18 kPa).^16^ This provided an ideal microenvironment, allowing organoids to survive for at least 15 weeks *in vitro* (Fig. 2a) without measurable changes in structure or size (Fig. 2b, Supp. Fig. 2a).

**Figure 2.**
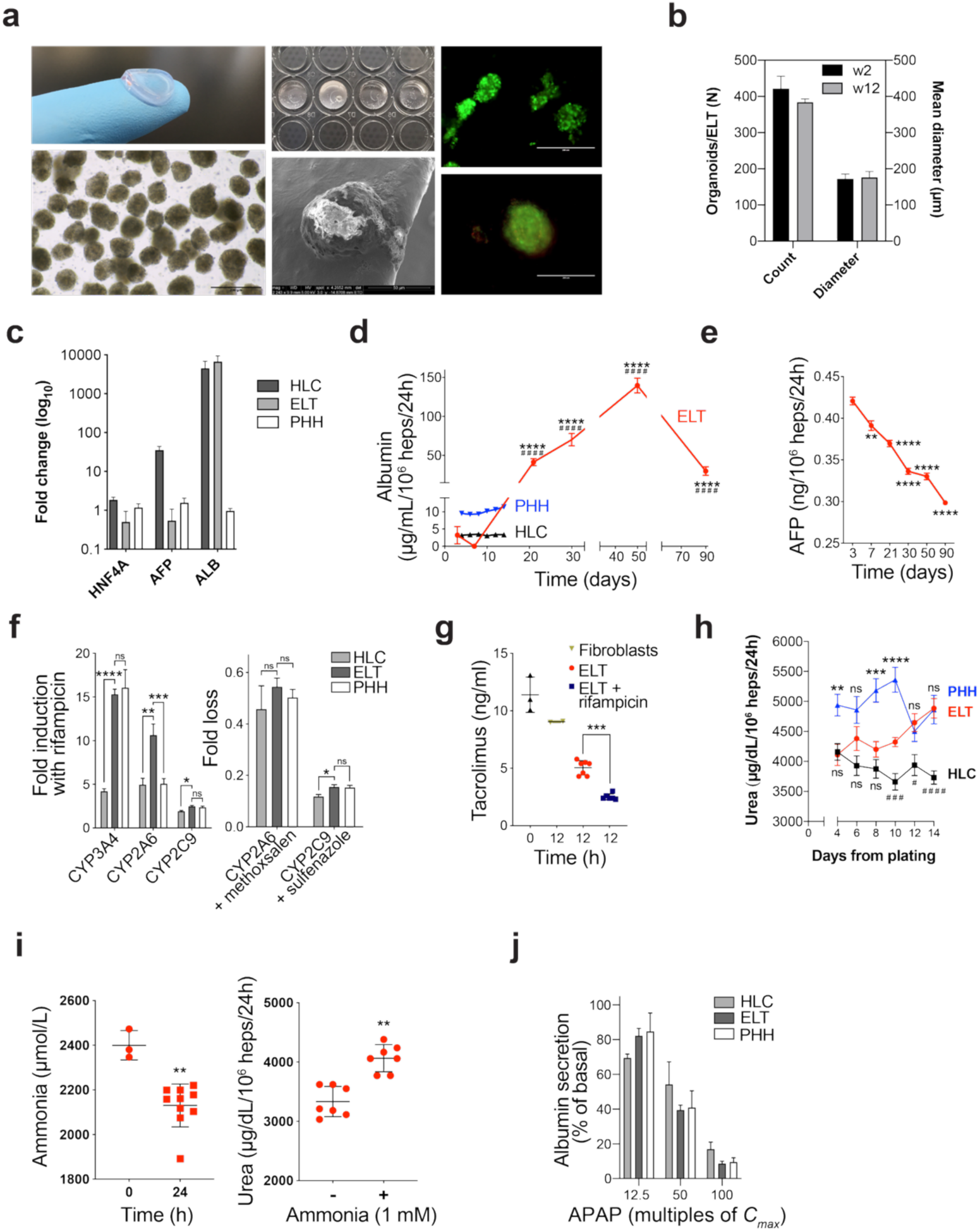
iPSC-derived ELT exhibits mature liver functions *in vitro*, matching or surpassing PHH. **(a)** ELT appearance (top left) with embedded liver organoids (bottom left: phase contrast, scale bar 200 µm. Top center: ELT used for *in vitro* assays. Bottom center: scanning electron microscope of an embedded organoid, scale bar 50 µm). Encapsulation provides an optimal environment for long-term survival of the organoids (top and bottom left: >90% live, fluorescent green calcein-AM-positive cells 4 weeks post-encapsulation, 5 independent experiments on 3 batches/biological replicates, representative images, scale bar 200 µm). **(b)** No significant growth or change in the number of organoids within the ELT was observed over time (average diameter and number or organoids/ELT at weeks 2 and 12 from encapsulation, w2 n=8 from 2 batches, w12 n=7 from 2 batches, *p*=not significant). **(c)** Expression of liver-specific genes in the ELT (2 weeks post-encapsulation) compared to PHH, iPSC-derived hepatoblasts and iPSC-derived HLC (Real-time RT-qPCR/TaqMan, values expressed as mean log10 fold change vs PHH ± SE; n=3 biological replicates for the ELT, n=6 biological replicates for HLC and PHH). **(d)** Albumin secretion from the ELT compared to iPSC-derived HLC and PHH (ELISA, mean ± SE normalized for 106 liver cells; n=28 ELT from 6 batches, n=12 PHH, n=10 HLC, unpaired *t* test with Welch’s correction, **** *p*<0.0001 vs. maximum value for PHH, #### *p*<0.0001 vs. maximum value for HLC). **(e)** Decreasing AFP secretion confirms ELT maturation into an adult phenotype (ELISA, mean ± SE normalized for 10^6^ liver cells; n=10 from 3 batches, paired *t* test vs. day 0). **(f)** The ELT showed inducible cytochrome P450 activity at least comparable to PHH (left; n=12 ELT from 2 batches, n=10 PHH, n=10 HLC, right; n=6 ELT from 2 batches, n=12 PHH, n=10 HLC; mean ± SE, Mann-Whitney U test). **(g)** Effective Cyp3A4-mediated tacrolimus metabolism by the ELT *in vitro* (liquid chromatography/mass spectrometry, n=7 ELT, n=3 T0, n=2 fibroblasts; Mann-Whitney U test). **(h)** The ELT synthesized urea more effectively than HLC, reaching levels comparable to PHH 2 weeks post-encapsulation (ELT n=38 from 6 batches/biological replicates, HLC n=10, PHH n=12; Mann-Whitney test, ** p<0.01, *** p<0.001, **** p<0.0001 vs PHH, # p<0.05, ### p<0.001, #### p<0.0001 vs HLC). **(i)** The ELT efficiently metabolized ammonia (left, 0.99 µmol/min/10^6^ liver cells; n=3 T0, n=10 from 2 batches T=24h, Mann-Whitney U test) into urea *in vitro* (right, n=7 from 2 batches). **(j)** ELT response to acetaminophen hepatotoxicity is comparable to PHH (% of basal albumin secretion, ELISA, mean ± SE normalized for 10^6^ liver cells, APAP dose expressed as multiples of *C*max; n=3). (* *p*<0.05, ** *p*<0.01, *** *p*<0.001, **** *p*<0.0001).

### The ELT achieves functional maturity *in vitro*

Liver organoids pursued their maturation within the optimal microenvironment provided by the ELT. By two weeks post-encapsulation, our engineered liver tissue expressed mature liver markers (Fig. 2c, Supp. Fig. 2b) and performed mature liver functions at levels equal to or exceeding PHH (Fig. 2d–h, Supp. Fig. 2c). These functions persisted for more than 12 weeks in culture. The ELT secreted liver-specific proteins at levels comparable to the human liver, including albumin, which reached values ∼13-fold higher than PHH and ∼40-fold higher than iPSC-derived HLC (p<0.0001), and maintained clinically relevant output levels for three months (Fig. 2d, Supp. Fig. 2b,c). In parallel, alpha-fetoprotein secretion progressively declined (Fig. 2e, Supp. Fig. 2b), confirming the acquisition of an adult phenotype by the hepatocytes. The ELT also demonstrated fully inducible and inhibitable cytochrome P450 (CYP) enzyme activity at levels comparable to PHH (Fig. 2f), including the capacity to metabolize tacrolimus effectively (Fig. 2g). But unlike PHH, the ELT’s functions remained stable upon shipping at 4°C and cryopreservation (Supp. Fig. 2d,e), which will facilitate the logistics of its use as a cell therapy product. The ELT synthesized urea at levels comparable to PHH (Fig. 2h) and was highly efficient at ammonia detoxification (Fig. 2i), a critical function for addressing hepatic encephalopathy in ALF. The extrapolated maximal *in vitro* ammonia clearance (86 µmol/min/g of liver tissue/L) was nearly 10-fold higher than basal human liver clearance capacity and surpassed 20% of the theoretical maximal capacity.^17^ In a proof-of-concept assessment of *in vitro* drug hepatotoxicity, the ELT behaved like PHH, but with less variability (Fig. 2j).

### Immune shielding enables off-the-shelf allogeneic therapy

Considered the condition’s narrow therapeutic window, a viable cell therapy for ALF necessitates a ready-to-use allogeneic product. To enable allogeneic administration of the ELT without the need for immunosuppression, we designed a delivery strategy that effectively shields the liver organoids from the host immune system. PEG-VS-based encapsulation provides long-term immune isolation, as the non-degradable material resists enzymatic degradation following implantation.^18,19^ The mesh size of 4% PEG-VS was determined by equilibrium swelling to be 23.4±0.5 nm (Fig. 3a) and remained unchanged following cryopreservation (Supp. Fig. 3a). This was confirmed by electron microscopy (Fig. 3b and Supp. Fig. 3b).^20^ Such a mesh size permits the diffusion of molecules, nutrients and proteins up to the size of albumin (∼66 kDa), while substantially restricting the passage of larger macromolecules such as immunoglobulins (∼150 kDa; Fig. 3c), and preventing any contact with the host immune cells.^21^

**Figure 3.**
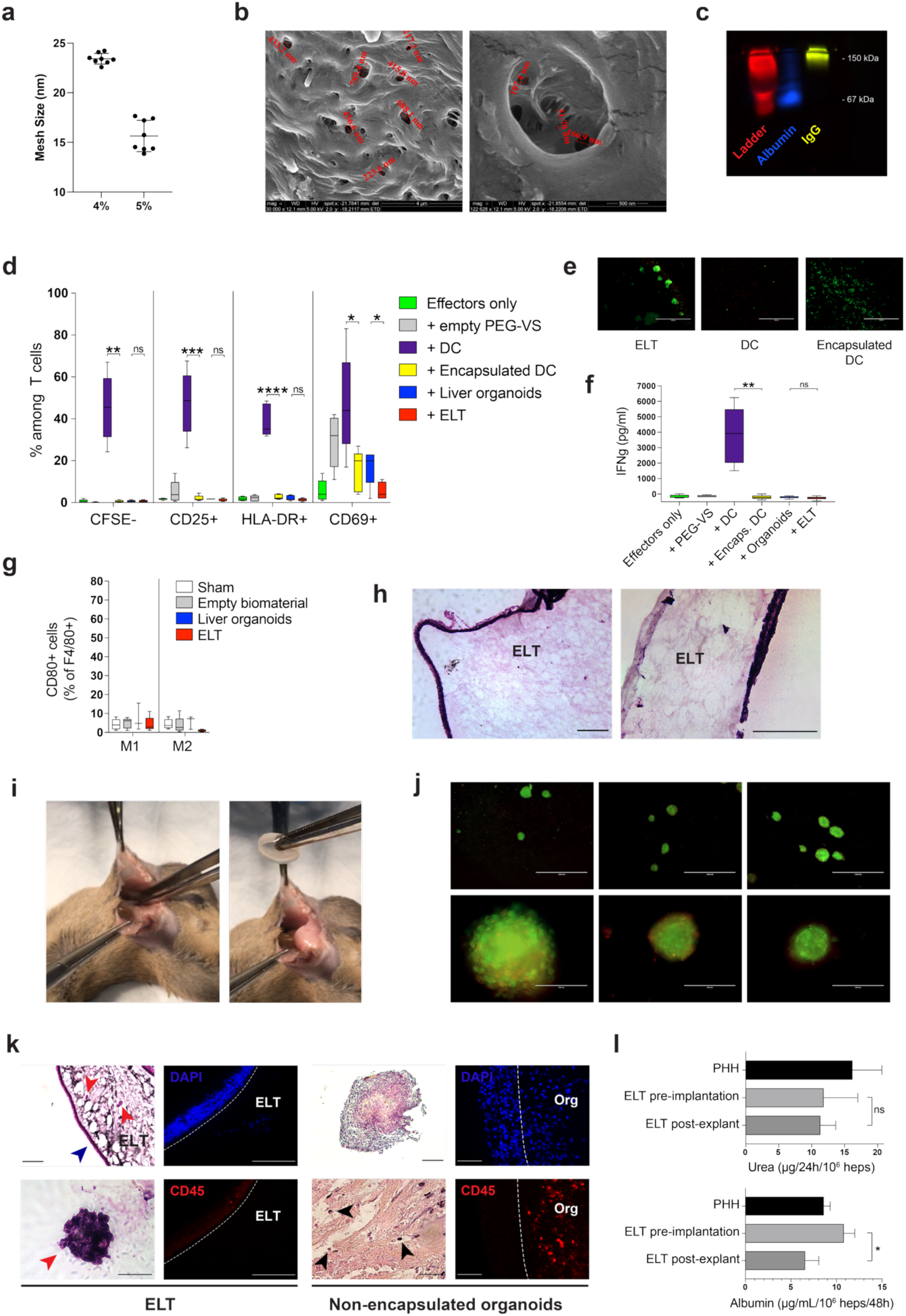
Encapsulation protects the ELT against rejection. **(a)** The mesh size of 4% and 5% PEG-VS calculated by equilibrium swelling (n=8, Mann-Whitney test, *p*<0.001). **(b)** Nanopores within crosslinked 4% PEG-VS ranged from approximately 20 to 700 nm (larger at the surface and decreasing deeper within the gel, scanning electron microscopy, scale bars: left 4 µm, right 500 nm). **(c)** Size exclusion: the porosity of the biomaterial allowed albumin (6 nm in radius) to diffuse within the hydrogel but prevented IgG molecules (20 nm) from penetrating (electrophoresis on 4% PEG-VS gel, n=3, representative image). **(d)** Encapsulation provided complete immune isolation *in vitro*, making liver organoid and activated dendritic cells (DC) invisible to allogeneic T cells (mixed lymphocyte reaction [MLR], % of T cells by flow cytometry; T cell proliferation measured by carboxyfluorescein succinimidyl ester staining [CFSE]; n=5 experiments with peripheral blood mononuclear cells from 5 donors; center line = median, box limits = upper and lower quartiles, whiskers = minimum and maximum; unpaired *t* test with Welch’s correction, * *p*<0.05, *\*\* p*<0.01, *** *p*<0.001. Details in Supp. Fig. 3c). **(e)** Encapsulation protected embedded cells from allogeneic T cell-mediated cytotoxicity (live/death assay, samples used for MLR showed in **d**, representative images; green=alive, red=dead). **(f)** Cytotoxic CD8+ T cells did not produce detectable IFN-**γ** when co-cultured with encapsulated allogeneic DC or the ELT (ELISA, n=5; center line = median, box limits = upper and lower quartiles, whiskers = minimum and maximum; unpaired *t* test with Welch’s correction, *\*\* p*<0.01); **(g)** The human ELT did not trigger significant peritoneal inflammation (flow cytometry analysis of M1 and M2 peritoneal macrophages 7 days post-implantation into immunocompetent mice; n=5, *p*=not significant). **(h)** Minimal foreign body reaction was noted when the 4% PEG-VS delivery device was implanted for 2 months into the peritoneal cavity of immunocompetent mice (representative images of the biomaterial explanted after 60 days within the peritoneal cavity of C57BL/6 mice; H&E, scale bar 200 µm). **(i)** Absence of adhesions or inflammation at explant of the ELT, 4 weeks post-implantation into immunocompetent mice, without immunosuppression (n=12, representative images). **(j)** Within the explanted ELT, human liver organoids were alive, with no measurable change in size or number (live/death assay; n=12 from 3 batches, representative images). **(k)** The explanted ELT (left panels) showed well-preserved liver organoids (red arrowheads, hematoxylin and eosin [H&E] staining) and no rejection. A thin fibrotic capsule containing some immune cells formed around the ELT (blue arrowhead and right images), but no immune cell infiltrated the biomaterial (immunostaining for CD45 and DAPI). Scale bars 200 µm except for 50 µm at bottom left (n=8 from 3 batches, representative images). In non-encapsulated human organoids used as controls (right panels), immune cell infiltration (black arrowheads) and necrosis of the liver tissue was found (left: H&E; right: immunofluorescence for CD45 and DAPI; scale bars 50 µm except for 200 µm at top left; n=5 from 2 batches, representative images). **(l)** The ELT explanted after 4 weeks were still functional and comparable to PHH, with foreign body reaction having only a negligible impact (top: urea secretion, n=9 ELT from 3 batches pre-implantation, n=4 ELT post-explant, n=4 PHH; *p*=not significant; bottom: albumin secretion, n=6 ELT from 3 batches pre-implantation, n=4 ELT post-explant, n=3 PHH; unpaired *t* test with Welch’s correction, * *p*<0.05).

We first evaluated the immune-isolating properties of this encapsulation *in vitro* using a mixed lymphocyte reaction. The ELT did not elicit T-cell activation, T-cell proliferation, or T-cell– mediated cytotoxicity, nor did CD8+ T cells secrete interferon-γ (IFN-γ) (Fig. 3d–f and Supp. Fig 3c). In contrast, non-encapsulated organoids induced mild T-cell activation, while activated dendritic cells (DC) triggered a strong response. When encapsulated within our system, these activated DCs no longer induced T-cell activation or proliferation, nor did they undergo lysis themselves (Fig. 3d-f).

The ELT was conceived to be implanted into the peritoneal cavity, without the need of any anastomosis, and perfused only by the peritoneal fluid, which in turn is in balance with the blood.^22,23^ By design, the non-degradable encapsulation prevented any vascularization of the ELT. The liver tissue physiologically operates at lower oxygen saturations than other abdominal organs because of its predominantly portal vascularization.^24,25^ This allowed for well oxygenated peritoneal fluid perfusing the biomaterial to meet the oxygen and nutrient requirements of the liver organoids, resulting in their long-term survival and function.^22,25,26^ When implanted into the peritoneal cavity of immunocompetent mice, without immunosuppression, the human ELT triggered no measurable peritoneal inflammation (Fig. 3g). Foreign body response against PEG-VS alone was minimal, forming only a thin fibrotic capsule around the implant after 8 weeks (Fig. 3h). Four weeks post-implantation, no vascular infiltration of the biomaterial was observed. The ELTs could be easily explanted with no noticeable adhesions or inflammation (Fig. 3i). Explanted ELTs harbored viable human organoids without immune cell infiltration (Fig. 3j,k), unlike non-encapsulated organoids, which were rapidly infiltrated and destroyed (Fig. 3k). Double encapsulation with a second thin layer of PEG-VS was needed to avoid that rare organoids might occasionally remain exposed on the surface of the ELT, where they were attacked by the recipient immune system (Supp. Fig. 3d). ELTs explanted after 4 weeks still performed liver functions at levels comparable to PHH (Fig. 3l). Such *in vivo* data confirmed the ELT’s biocompatibility and demonstrated that encapsulation within PEG-VS prevented any contact between the recipient’s immune system and the embedded organoids, with no rejection despite the implantation of a human cellular product in immunocompetent mice.

### Encapsulation protects against tumor formation

The ELT was designed as a fully contained system. In addition to preventing immune reactions, encapsulation enhances safety by restricting dissemination of embedded cells, thereby reducing the risk of tumorigenicity, a key concern for iPSC-based therapies. When implanted into immunodeficient NOD scid gamma (NSG) mice, the ELT did not produce tumors after 16 weeks, whereas unencapsulated undifferentiated iPSC formed teratomas in these mice after 6-8 weeks (Supp. Fig. 4a). Although no residual undifferentiated iPSCs were detected among the cells composing our organoids,^7^ encapsulation was stringent enough to eliminate any theoretical risk of teratoma formation, as further shown by encapsulated undifferentiated iPSC generating no ectopic growth 16 weeks post-implantation in NSG mice (Fig. 4a). To further assess the protective effect of encapsulation against tumorigenicity, we transformed iPSC-derived hepatoblasts into liver cancer cells capable of generating undifferentiated hepatoblastomas upon injection into NSG mice (Supp. Fig. 4b). When these liver cancer cells were encapsulated into PEG-VS hydrogels (Fig. 4b), no tumor were observed 16 weeks post-implantation (Fig. 4c-e, Supp. Fig. 4c), confirming the safety of our encapsulation approach.

**Figure 4.**
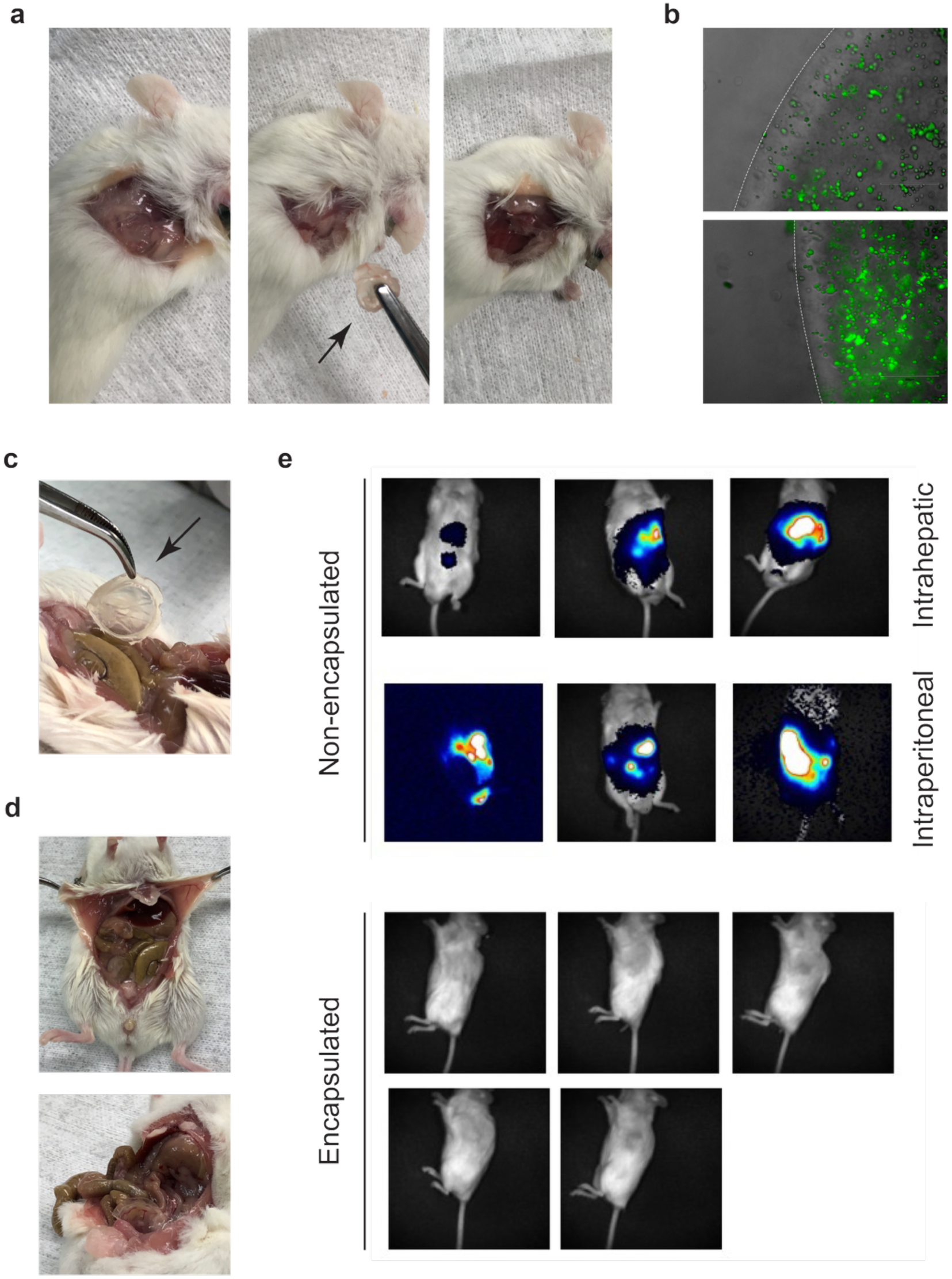
The ELT is a fully contained system designed to minimize risks of tumorigenicity. **(a)** Encapsulation prevented teratoma formation by avoiding the spread of embedded cells: implantation of encapsulated undifferentiated iPSC into immunodeficient NSG mice resulted in no ectopic tissue found after 16 weeks (n=4 biological replicates, representative images; controls in Supp. Fig. 4a). **(b)** Encapsulated GFP/luciferase-tagged undifferentiated hepatoblastomas cells derived by the transformation of iPSC-derived hepatoblasts (representative image of 7 biological replicates, scale bar 300 µm). **(c-e)** Encapsulation protected NSG mice against the development of malignant liver cancers 16 weeks after the implantation of the GFP/luciferase-tagged undifferentiated hepatoblastomas cells: **c)** intact encapsulation at explant; **d)** absence of tumor formation at necropsy (representative images); **e)** live imaging showing absence of tumor formation with encapsulation compared to significant luciferase-positive masses developing in controls (n=7 biological replicates; histology of the tumors showed in Supp. Fig. 4b, additional images maximizing sensitivity showed in Supp. Fig. 4c).

### ELT as a treatment for acute liver failure

Once the safety of the ELT and its functional survival into the abdominal cavity established, we assessed its efficacy in a clinically relevant mouse model of ALF (Fig. 5a). Immunocompetent mice were injected with a lethal dose of CCl₄, and by 24 hours, they showed marked liver injury, hyperammonemia, and neurological signs compatible with HE.^27,28^ Animals were then treated either with the human ELT or an empty PEG-VS (control), which were implanted into the peritoneal cavity, the two groups being homogeneous for ALF severity (Fig. 5b). No immunosuppression was administered. At 5 weeks, 68% of mice receiving ELTs survived, compared to 29.2% of controls (p=0.0098, hazard ratio 0.37; Fig. 5c). All deaths were associated with severe signs of HE. Plasma ammonia levels 48h post-implantation decreased in mice receiving the ELT compared to controls (Fig. 5d). ELT-treated mice showed reduced incidence and severity of neurological impairment (Fig. 5e-g) and a higher recovery rate even among those presenting more severe HE (Fig. 5h). After 5 weeks, the ELT was then explanted from surviving animals and mice were followed for 2 additional months (Fig. 5i). The explanted ELT showed no signs of rejection. At day 90 post-treatment, all animals were alive and healthy, with normal ALT levels and liver histology (Supp. Fig. 5a). This outcome suggests that the ELT provided essential bridging functions by supporting vital synthetic and metabolic processes and by managing complications such as HE while the host liver recovered. The implant could then be safely explanted, leaving the animals cured.

**Figure 5.**
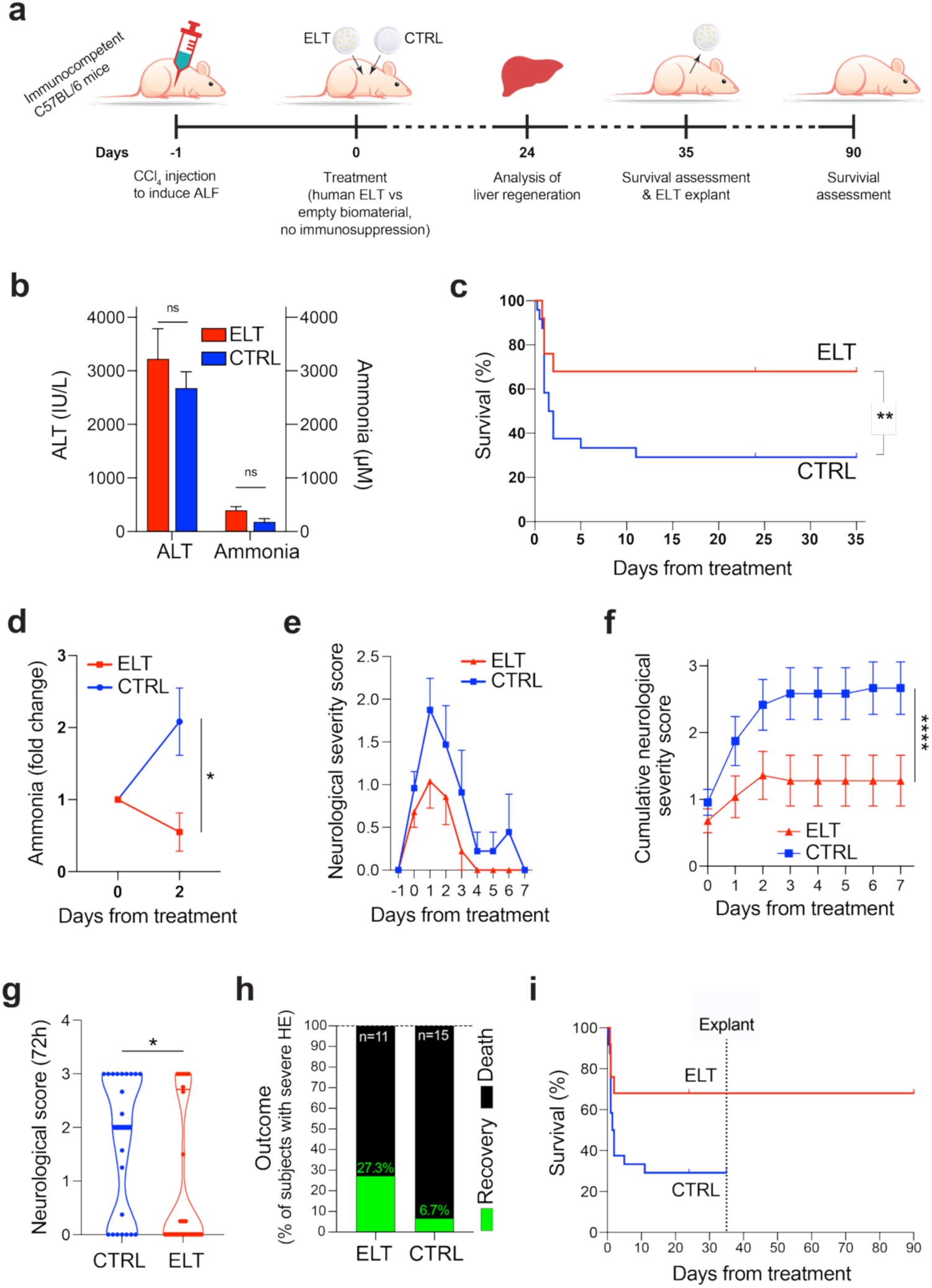
Human ELTs effectively treats ALF in immunocompetent mice without immunosuppression. **(a)** Experimental design. **(b)** Twenty-four hours after CCl4 injection the mice showed biochemical signs of severe ALF (elevated plasma alanine aminotransferase [ALT] levels, n=7 per group, and hyperammonemia, n=4 per group), with no difference between animals randomized to receive the ELT and those to receive the empty biomaterial (*p*=ns). **(c)** The ELT improved survival of the treated animals (68% vs. 29.2% in controls, ** p<0.01; n=25 animals with ELT from 3 batches, n=24 animals CTRL, logrank test). **(d)** Ammonia levels 48h after treatment decreased in mice receiving the ELT and increased in controls (fold change vs. before treatment, n=3 ELT, n=4 CTRL, Welch’s t test, * p<0.05). **(e)** ELTs reduced the incidence and severity of HE in mice with ALF (neurological and behavioral score based on a modified Irwin test; n=25 ELT, n=24 CTRL; mixed-effects model/REML, *p*<0.05). **(f)** Cumulative neurological was lower in mice receiving the ELT than in CTRL, as a result of the effect of the ELT on HE (the score taks into account death/sacrifice following coma over the first 7 days from treatment; n=25 ELT, n=24 CTRL, 2way ANOVA, *p*<0.0001). **(g)** Neurological and behavioral score over the first 3 days from treatment was significantly lower for mice treated with the ELT than for controls (n=25 animals for ELT, n=24 CTRL, unpaired t test with Welch’s correction, * *p*<0.05). **(h)** Higher recovery rate in ELT-treated mice with severe early HE compared to controls (n=11 ELT, n=15 CTRL). **(i)** ELTs were safely explanted at day 35 and all mice survived with a regenerated liver (histological images showed in Supp. Fig. 5). (Controls were sacrificed at day 35 for ethical reasons.)

We also examined liver regeneration by assessing a subset of survivors at day 24 post-implantation (Fig. 5c), when ALT levels showed no active liver injury in mice having received the ELT and only mild ALT elevation in the control group (Fig. 6a). Livers of mice that received ELTs appeared normal at examination (Fig. 6b), with negligible residual liver damage (Fig. 6c), whereas surviving controls still exhibited hepatomegaly, centrilobular inflammation, and apoptosis. Improved liver regeneration was already observed in treated animals 72h after ELT implantation (Fig. 6d-e), suggesting an effect of the ELT in accelerating the regeneration of the recipient’s own liver.

**Figure 6.**
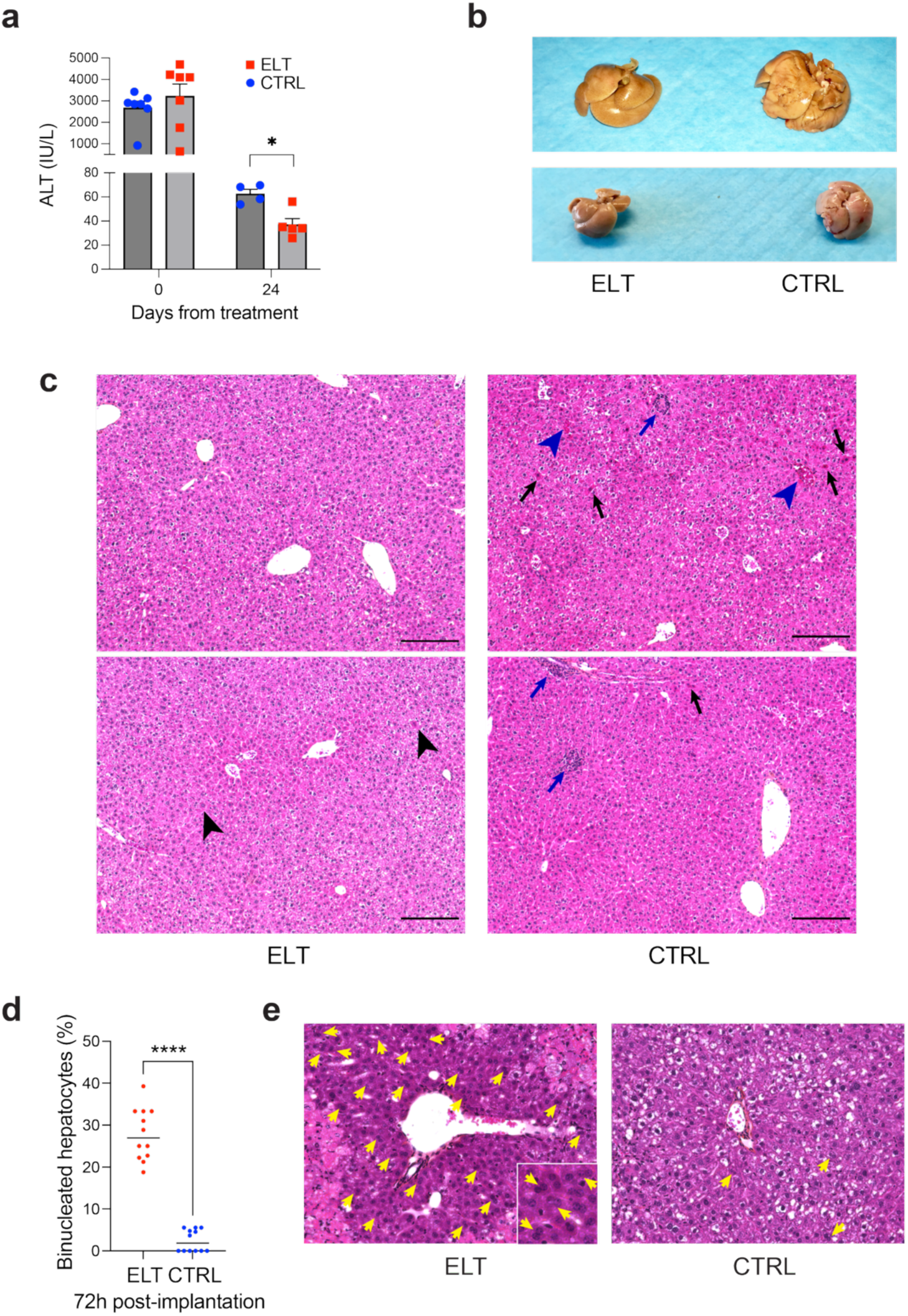
The ELT promotes the regeneration of the recipient’s own liver. **(a)** At day 24 post-treatment of mice with CCl4-induced ALF, survivors having received the ELT had normalized ALT levels (n=5), whereas surviving controls showed a significantly reduced (n=4), although not yet normal levels (34±13 vs 63±27 IU/L, Mann-Whitney, * *p*<0.05). **(b)** ELT promotes liver regeneration: at day 24 post-implantation (see Fig. 5a), the liver of mice having received the ELT had a normal size and appearance compared to surviving controls’, which were still enlarged and softer (n=5 ELT, n=2 CTRL, representative images of 2 animals per group). **(c)** Microscopic appearance of livers from the ELT group at day 24 (left) were mostly normal, with only rare hepatocytes with condensed cytoplasm in zone 3 (black arrowheads) and no inflammation, necrosis or apoptosis, whereas controls showed centrilobular venous congestion (blue arrowheads), lobular inflammation (blue arrows) and rare acidophilic bodies (back arrows; H&E, scale bar 200 µm; n=5 ELT, n=2 CTRL, representative images of 2 animals per group). **(d,e)** The ELT accelerated host liver regeneration: 72h after implantation, the number of binucleated hepatocytes was significantly higher in animals receiving the ELT than in controls (left: n=4, 3 measurements per animal, p<0.0001; right: representative images, H&E staining, some binucleated hepatocytes are identified by yellow arrowheads, insert shows magnified area).

## DISCUSSION

Survival with their native liver being poor, people with ALF need to be transplanted within days from the diagnosis, before their disease becomes irreversibly too severe. But liver transplantation still entails significant short-term mortality and morbidity risks, requires lifelong immunosuppression and is often complicated by severe long-term hepatic and extra-hepatic problems. Such risks and long-lasting complications seem disproportioned for many patients with ALF who would just need temporary replacement of liver functions while their own liver regenerates. The data showed above suggest that our engineered liver tissue has the potential to be developed into a safe and effective allogeneic cell therapy for ALF, improving survival, alleviating severe complications such as hepatic encephalopathy, and promoting liver regeneration, all without the need for immunosuppression. Once the patient’s own liver regenerated, the ELT is easily explanted, potentially leaving the person cured, with no residual foreign cell in the body nor need for long-lasting treatment.

Despite almost 30 years of clinical experience, liver cell transplantation was not able to show convincing efficacy in ALF.^5,29,30^ Limited by donor shortage and high functional variability, intraportal administration of primary liver cells provides a short-lived mass effect, followed by a slow and very limited engraftment (see below). Intraperitoneal administration of encapsulated hepatocytes showed clinical promise for ALF, but does suffer from many of the same limitations.^31^ Compared to primary liver cells, the ELT has the added advantage of superior and longer-lasting functionality, consistency, better stability upon cryopreservation, and derivation from unlimited source material. Several elegant stem cell-derived approaches have been described, but require engraftment, in vivo maturation and immunosuppression, with still uncertain long-term safety profiles.^9,11,27,32,33^ Extracorporeal approaches to liver function replacement have historically disappointed, although preclinical promise was shown using porcine hepatocytes.^34–37^ Immediately functional upon implantation, designed as a transient, fully contained implant, and not requiring animal sacrifice, engraftment or immunosuppression, the ELT presents many advantages over these approaches.

The design of the ELT was guided by patient needs, with particular attention to safety considerations and clinical feasibility. An off-the-shelf design is strictly required to allow for rapid deployment to patients within the ALF’s very short therapeutic window. This implies an allogeneic approach and the capacity to effectively cryopreserve the product for easy storage and shipment. The use of a single iPSC line as starting material allows for highly consistent manufacturing from a virtually unlimited source. As shown above, thanks to the recreation of the liver microenvironment within the encapsulating biomaterial, the ELT achieves maturation during the manufacturing process, which allows it to be immediately effective in replacing liver functions upon implantation, and makes it uniquely suited for even the most acute forms of liver failure.

Implantation of the ELT consists in a short and minimally invasive procedure that does not require any complex anastomosis. The ELT does not need engraftment nor any vascularization since all the exchanges are accomplished through perfusion by the peritoneal fluid, which in turn interacts with the highly vascularized peritoneum and is in balance with the blood.^22,26^ Such an approach is supported by extensive clinical experience with peritoneal dialysis to treat hyperammonemia,^38–41^ and further confirmed by initial clinical reports using encapsulated hepatocytes.^31^ Engraftment has constituted one of the major hurdles for liver cell transplantation, with the reticuloendothelial system causing rapid cell loss within a few hours from intraportal administration, despite immunosuppression.^5,42,43^ Our data showed that the nondegradable biomaterial used in the ELT is fully biocompatible and assures protection from the host’s immune system, all while allowing for optimal perfusion and flow of toxic compounds, drugs to be metabolized, and synthesized proteins to be excreted. Unlike pancreatic islet, which have high oxygen requirements,^44^ hepatocytes within the liver are mostly perfused by venous blood coming from the portal vein, with much lower oxygen requirements.^24,25^ Although confirmation in more representative large animal model will be needed, foreign body reaction appeared negligible in a rodent model that was previously validated to assess fibrosis upon intraperitoneal implantation.^45^ If fibrotic reaction is likely to become limiting to exchanges between the ELT and the peritoneal fluid/peritoneum/blood over time, it does not seem to have any meaningful impact over the few weeks needed for functional replacement before sufficient liver regeneration is achieved. The ELT is indeed designed as a transient implant, which is removed after 3-5 weeks, once vital liver functions are restored to the point to support the patient’s life.^46,47^ Because of this and of the specific perfusion needs of liver organoids, we do not expect the ELT to suffer from the same problems that have hampered the clinical development of encapsulated cell therapies for diabetes mellitus.^48^

Immune shielding via macroencapsulation within PEG-VS prevents rejection and eliminates the need for immunosuppressing patients that are already at very high risk of infection.^49^ Rejection has hampered the progress of hepatocyte transplantation and other promising liver cell therapies.^5,27,42,43,50^ As previously described for other tissues, the tight size exclusion provided by the biomaterial’s controlled porosity protects the liver organoids from both T and B cell-mediated responses, as seen in allogeneic and xenogeneic contexts shown above. This enables the allogeneic approach, which is not only required to treat such an acute condition, but also to make the therapeutic product viable from a manufacturing, logistic and commercial point of view.

The ELT was designed to address key safety concerns that constitute a challenge for other stem cell-based approaches, among which the theoretical risk of teratoma or tumor formation intrinsic to the use of pluripotent stem cells as source material.^51,52^ We previously showed that the cells composing our liver organoids are terminally differentiated, with no residual undifferentiated stem cells.^7^ We demonstrated here that the ELT’s encapsulation prevents any dissemination of implanted cells throughout the recipient’s body. If any cells were to escape, their allogenic nature and the absence of immunosuppression would ensure their rapid clearance by the host immune system. Together with the transient design of implantation, these features provide a compelling safety profile.

As mentioned above, unlike most stem cell–derived liver products,^11–13,53^ the ELT achieves advanced maturation *in vitro*, and can be cryopreserved once mature, enabling rapid deployment to patients with ALF, at any hospital worldwide, within the disease’s narrow therapeutic window. Although differentiation protocols evolved over the years, hepatocyte-like cells derived from stem cells do not achieve full maturation in vitro.^7,8,54^ Further lengthy maturation in vivo is thus required, which prevents them from being used for ALF.^9^ The addition of non-parenchymal liver cells into a 3-dimensional structure allows for the recreation of cell-to-cell contacts and gradients of paracrine signals that constitute the liver microenvironment.^7,10,11,55^ This rapidly improves the maturation of the hepatic progenitor cells within the organoids, although not enough to be effective in the most severe forms of ALF.^13,32^ In contrast, thanks to the encapsulation into a biomaterial recreating the physical properties of the adult liver, our organoids achieve full maturity for many vital liver functions *in vitro*. With diffusion of nutrients and oxygen optimized by the controlled size of the organoids and encapsulation design, these functions are then maintained for several weeks, unlike PHH that rapidly de-differentiate upon digestion from the liver.^56,57^ And unlike PHH, whose quality unpredictably varies between donors,^58^ ELTs show negligible variability in liver functions between batches. This is due to the off-the-shelf approach (i.e. the use of a single iPSC line) and the robustness of the differentiation protocol,^7^ which was designed to ease scalability and transition to a cGMP manufacturing environment, in order to streamline clinical translation.

We also showed above that, thanks to the controlled composition and size of the organoids, as well as their encapsulation, the ELT can withstand cryopreservation without any meaningful functional loss. This significantly differentiates the ELT from PHH, which notoriously suffer from cryodamage.^59^ Such a feature is of outmost importance to allow for the easy storage, shipment and deployment of the therapeutic product to the patients within the therapeutic window.

The number of cells needed to provide sufficient functional replacement in ALF is unknown. In case of liver transplantation, a graft of less than 40%-50% of standard liver volume is associated with unfavorable outcome.^60^ Whereas 5%-10% of the liver mass was shown to be needed in case of metabolic liver diseases, no reference exists for ALF.^29,61^ Preclinical studies using bioartificial liver devices suggested that 10% of the total liver mass was needed to rescue ALF.^36^ Nevertheless, anecdotical clinical experience with PHH showed some efficacy with <1% of the liver mass.^31^ Although the dose tested in the efficacy proof of concept shown above was very low, representing approximately 0.5% of the theoretical number of hepatocytes in the mouse liver,^62^ it was sufficient to reverse ALF in most animals. Nevertheless, better efficacy can reasonably be expected with a higher dose, which will be targeted upon scale-up of the manufacturing process.

## Conclusion

Overall, the ELT showed promise as an off-the-shelf advanced therapy to treat ALF. It overcomes many of the challenges that have hampered previous attempts to develop effective cell therapies for liver disease, with a unique safety profile. Designed for future clinical translation, with feasibility, scalability and easiness of deployment in mind, it is based on robust differentiation protocols and delivery method, and a simple route of administration. Already transferred to the industry, the ELT is undergoing further process and preclinical development in preparation for clinical trials. Pending validation in large animal models and clinical trials, the ELT has the potential to replace liver transplantation for most patients with ALF, and be a bridge for transplant for the most severe cases.

## ONLINE METHODS

### Ethical approval

This project was approved by the CHU Sainte-Justine internal review board (CER protocols 1386 and 2195), and committee for good laboratory practices for animal research (CIBPAR protocol 655-2600) and by the Canadian Institute of Health Research’s Stem Cell Oversight Committee.

### iPSC generation and maintenance

Human iPSCs were reprogrammed from skin fibroblasts and PBMCs of 3 healthy donors by Sendai-virus delivery using the CytoTune-iPS 2.0 Sendai Reprogramming Kit (ThermoFisher Scientific), following the manufacturer’s instructions. The colonies with suggestive morphology were manually picked based on live staining for the pluripotency marker TRA-1-60 and then cultured as iPSC thereafter, with single colony sub-cloning for the first 5 passages. After passage 10, a temperature shift (incubation at 39°C for 72h) was performed to remove the cMyc gene, and the absence of Sendai virus was confirmed by RT-qPCR. PSC cultures were maintained on vitronectin-coated plates (ThermoFisher Scientific) in Essential 8 Flex medium (ThermoFischer Scientific) at 37°C in humidified atmosphere containing 5% CO_2_ and 4% O_2_.

### iPSC differentiation into hepatoblasts

iPSCs were differentiated into hepatoblasts following our previously described protocol.^7^ Hepatoblasts were consistently generated from all 3 iPSC populations (9 differentiations overall) with negligible variability. Briefly, the cells were dissociated by TrypLE (Life Technologies) to single cells and seeded on human recombinant laminin 521 (BioLamina)-coated plates in Essential 8 Flex medium supplemented with 1% RevitaCell (Life Technologies), at a density of 7 x 10^5^ cells/cm^2^. Differentiation was started (day 0) when the cells reached around 70% confluence by changing the medium to RPMI-B27 minus insulin (Life Technologies) supplemented with 1% knockout serum replacement (KOSR, Life Technologies). For the first 2 days, the cells were exposed to 100 ng/ml Activin A (R&D Systems) and 3 μM CHIR-99021 (Stem Cell Technologies), and then for the 3 following days to 100 ng/ml Activin A alone. Subsequently, RPMI-B27 (minus insulin) medium was supplemented with 20 ng/ml BMP4 (Peprotech), 5ng/ml bFGF (Peprotech), 4 μM IWP-2 (Tocris), and 1 μM A83-01 (Tocris) for 5 days, with daily medium change. At day 10, the medium was changed to RPMI-B27 (Life Technologies), supplemented with 2% KOSR, 20 ng/ml BMP4, 5 ng/ml bFGF, 20 ng/ml hepatocyte growth factor (HGF, Peprotech) and 3 μM CHIR-99021 for 5 days, with daily medium change. During all the differentiation process, the cells were kept at 37°C, ambient O_2_ and 5% CO_2_.

### iPSC differentiation into mesenchymal progenitor cells

iPSCs were differentiated into mesenchymal progenitor cells (MPC) as previously described, with modifications.^63^ Briefly, iPSCs were seeded after a single-cell passaging with TrypLE on human recombinant laminin 521-coated plates, and the differentiation was started when 50% confluence was reached. The medium was changed to DMEM high glucose with 10% KOSR for 2 weeks, with every-other-day medium change. After 2 weeks, the cells were passed on plastic plates (Corning) and, after 4 subsequent passages, they were considered as mesenchymal progenitor cells, and were fully characterized.

### iPSC differentiation into endothelial progenitor cells

iPSCs were differentiated into endothelial progenitor cells (EPC) as previously described, with modifications.^64^ Briefly, iPSCs were seeded after a single-cell passaging with TrypLE on human recombinant laminin 521-coated plates, and the differentiation was started when 60% confluence was achieved. On day 1, Essential 8 Flex medium was changed for Stemdiff APEL II medium (Stem Cell Technologies) supplemented with 6 μM CHIR-99021 and 25 ng/ml BMP4, for 2 days. On day 3, the medium was changed to Stemdiff APEL II supplemented with 25 ng/ml BMP4 and 50 ng/ml VEGF (Life Technologies), for 4 days, with daily medium change. After 4 days, the cells were passed on human recombinant vitronectin (Life Technologies)-coated plates in EBM2-EGM2 media (Lonza). After 1 week, a cell sorting was performed for the CD31+ CD144+ double positive cells.

### Generation of controlled-size liver organoids

iPSC-derived hepatoblasts (3×10^5^ per ELT for *in vitro* experiments and 1×10^6^ per ELT for *in vivo* studies), MPC and EPC (respectively 1:0.2:0.7 ratio) were resuspended in a composite medium (50% EBM2-EGM2, 50% HBM-HCM supplemented with 1% KOSR), supplemented with 10 μM dexamethasone, 20 ng/ml oncostatin M (R&D Systems), 20 ng/ml HGF (Peprotech), and seeded in ultralow-attachment microcavity plates (Corning Elplasia). Within the microcavities, the cells were maintained in 50% EBM2-EGM2, 50% HBM-HCM 1% KOSR, supplemented with 10 μM dexamethasone, 20 ng/ml OSM, 20 ng/ml HGF for 5 days with daily half medium change. The self-aggregated organoids were then harvested and encapsulated to form the ELT.

### Encapsulation and ELT generation

Liver organoids were resuspended in 4%(wt/vol), 4-arm, 20-kDa PEG-vinyl sulfone (PEG-VS, JenKen Technology) and 0.4% Irgacure 2959 (Sigma-Aldrich) in dPBS. N-vinyl-2-pyrrolidone (0.1% wt/vol, Sigma-Aldrich) was added to accelerate PEG-VS photo-crosslinking and minimize cytotoxicity.^18^ For each ELT, the obtained suspension was then casted in a well of a 96 well-plate (for *in vitro*) or a 48 well-plate (for *in vivo*). The biomaterial was then photo-crosslinked though UVA irradiation (365 nm, 5 minutes, 115.6 µW/cm^2^). The size of the molds used for casting was chosen to assure a final thickness of the ELT of 2 mm, for all uses. A second thin layer of biomaterial was then applied on the ELT and quickly photopolymerized. A double encapsulation was required to prevent the recipient’s immune cells to attack the rare organoids that might remain partially exposed on the surface of the ELT (Supp. Fig. 3d).

### Flow cytometry

Cells were harvested and aliquoted (0.5-1×10^6^ cells in each assay tube). They were stained with 100 μl of fluorochrome-conjugated primary antibody solution (membrane antigens) for 20 minutes at room temperature protected from light. Cells were subsequently fixed with 4% paraformaldehyde for 10 minutes at room temperature. In case of intracellular staining, cells were permeabilized with 1% Triton X-100 after washing, and then stained with 100 μl of fluorochrome-conjugated primary antibody solution for 20 minutes at room temperature, protected from light. After extensive washing, cells were resuspended in 0.5 ml staining buffer (1% PBS-BSA) and kept at 4°C until analysis. The used fluorochrome-conjugated primary antibodies were: FITC anti-human CD90 (BD Bioscience), PE anti-human CD73 (BD Bioscience), APC anti-human CD146 (BioLegend), PerCP-Cy5-5-A anti-human CD105 (BD Bioscience), PE anti-human CD20 (BioLegend), PE-Cy7-A anti-human HLA-DR (BioLegend), PE anti-human TRA-1-60 (BioLegend), PE anti-human SSEA4 (Stemcell Technologies), FITC anti-human CD34 (BD Bioscience), PE anti-human CD45 (BioLegend), PE anti-human CD144 (BD Bioscience), PE anti-human VEGFR2 (BioLegend), PE anti-human CD31 (BD Bioscience), FITC anti-human CD14 (BioLegend). We used the following gating strategy: 1) preliminary compensation with beads was carried out to minimize fluorescence spillover if multicolor staining was used; 2) scatter gating (gating on forward scatter [FSC] and side scatter [SSC]), excluding debris and background noise by setting the appropriate FSC threshold; 3) fluorescence gating using unstained cells. When multiple cell lines were analyzed by flow cytometry at the same time, we prepared unstained cells for each line. For mixed lymphocyte reaction analysis, after the gating strategy described above, alive cells were gated based on exclusion of 7AAD-positive cells and CD3-positive cells were subsequently selected. Data was acquired using BD FACSDiva software v6.1.3 on a BD FACSCanto II cell analyzer (BD Biosciences). Data analysis was performed with FCS express v4 (De Novo software).

### Gene expression analysis

Total RNA was extracted using ReliaPrep RNA Miniprep system (Promega) from cultured cells and ELT and used as a template for synthesis of single-strained cDNA. Reverse transcription was performed to obtain cDNA (Omniscript, Qiagen). The PCR reaction mixes were prepared and afterwards loaded in the plates. The plates were sealed, centrifuged and then loaded in the instrument (Life Technologies). Standard TaqMan qPCR reaction conditions were used. The following TaqMan gene expression assays (Thermo Fisher) were used: Hs99999905_M1 (GAPDH), Hs00609411_M1 (Albumin), Hs002230853_M1 (HNF4A), Hs00173490_M1 (AFP). Data were analyzed using the comparative CT (Λ1Λ1CT) method for calculating relative quantitation of gene expression and expressed as log_10_ fold change.

### Animals and *in vivo* experiments

Animal experiments were approved by CHU Sainte-Justine committee for good laboratory practices for animal research (CIBPAR protocol 655-2600) and conducted in accordance with the Canadian Council on Animal Care (CCAC) guidelines in CHU Sainte-Justine’s CCAC-certified animal core facility. The mice were bred and maintained according to the CHU Sainte-Justine institutional guidelines for the use of laboratory animals. The number of animals for each experiment was calculated based on the results of pilot studies or previously published literature. *In vivo* efficacy: Fifty-four female C57BL/6 mice aged 6-8 weeks, with *ad libitum* access to appropriate standard diet, were injected intraperitoneally with carbon tetrachloride (CCl_4_, 3 ml/Kg of body weight vehiculated in mineral oil). Three mice died within the first 24 hours. Twenty-four hours after the administration of the CCl_4_ (day 0), blood was taken from the mice to measure liver enzymes and ammonia levels in order to confirm liver failure. Animals were randomized to receive either receive the ELT or the empty biomaterial. Subsequently, under anesthesia with isoflurane, the ELT was implanted by laparotomy in the abdominal cavity of 27 mice (50,000 liver cells/g of body weight; treated group), while 24 mice received the empty biomaterial (control group). Two mice from the ELT group did not wake up after surgery. Only animals surviving the surgery were included in subsequent analysis. Blood tests were repeated at day 2, 9, and 16 to monitor liver enzymes and ammonia. Mice were monitored each 2 hours for the first 48 hours after the surgery and then twice a day, with neurological and behavioral evaluation performed according to a modified Irwin test.^65^ The following clinical parameters were enregistered and graded accordingly: body position (normal = 0, sitting crouched = 2, lying on one side = 3), locomotor activity (normal = 0, restless = 1, reduced exploration = 1, no activity or movement = 2), whole body tremor (absent = 0, present = 1), ataxia (absent = 0, present = 2), reactivity to touch (normal = 0, reduced = 1, absent = 2). The neurological score was calculated at fixed timepoint adding the scores listed above. A coma score was also calculated at every assessment (movements, reflexes, painful response to tail sting, posture, and strength), as previously described.^66^ All the mice that have reached a grade 4 in the coma scale were euthanized. At day 24, 5 mice form the treated group and 2 mice from the control group were sacrificed to assess liver regeneration. At day 35 all surviving controls were sacrificed, while all surviving mice from the treated group underwent the explant of the ELT under anesthesia and were then followed up until day 90, and then euthanized. *In vivo safety:* To assess peritoneal inflammation, 20 C57BL/6 mice (6-8 weeks of age), with *ad libitum* access to appropriate standard diet, received either the ELT, the empty biomaterial, non-encapsulated liver organoids, and 0.9% sodium chloride (5 mice per group) into the peritoneal cavity (standard laparotomy), without any immunosuppression. After 7 days the mice were sacrificed, and the peritoneal liquid was harvested by washing the peritoneal cavity to assess the presence of macrophages. In order to evaluate foreign body reaction and rejection, 12 C57BL/6 mice aged 6-8 weeks received the ELT (implanted into the peritoneal cavity through standard laparotomy), without any immunosuppression. After 4 weeks, the mice were sacrificed and examined to assess for signs of adhesions and inflammation. The ELT were recovered from the peritoneal cavity and fixed for histological analysis to evaluate immune cell infiltration (N=8) or put back in culture to assess cell viability and liver-specific functions (N=4). In order to assess the tumorigenicity of the ELT, 8 NSG mice received either free undifferentiated iPSC or encapsulated undifferentiated iPSC (4 mice per group) subcutaneously. The mice that received free iPSC were sacrificed after 8 weeks because of the evidence of the formation of big tumors (confirmed to be teratomas at histological analysis), while mice having received encapsulated iPSC were sacrificed after 16 weeks and thoroughly examined to look for ectopic tissue formation. Six NSG mice received an intrahepatic injection of 500,000 non-encapsulated luciferase-tagged iPSC-derived, transformed hepatoblasts, while 7 NSG mice received 500,000 macroencapsulated luciferase-tagged liver cancer cells into the peritoneum. Briefly, mice were anesthetized, and a 15 mm incision was made underneath the ribcage on the left ventral flank. Using a glass capillary, a 10 µL injection was made at about 2 mm depth in the liver lobe. Incisions were sutured, and the animal was treated with buprenorphine daily for 2 days following procedure. Mice that received non-encapsulated luciferase-tagged liver cancer cells were sacrificed after 4-6 weeks because of the evidence of the formation of liver tumors at live imaging. The mice that received encapsulated luciferase-tagged liver cancer cells were sacrificed after 16 weeks and examined to assess for signs of tumor formation. *In vivo* monitoring of tumor growth was done at regular intervals by bioluminescence tracking of firefly luciferase-expressing tumor cells. To do so, a 30 mg/mL solution of D-luciferin (PerkinElmer 122799) was injected i.p. at a dose of 150 mg/kg and imaged after 10 minutes without filters on the Q-Lumi *In Vivo* Imaging System (MediLumine, Montreal). Signal normalization and analysis was done automatically for all time points using FIJI macros and expressed in radiance (photons · s^-1^ · sr^-1^ · cm^-2^) integrated density (Area · mean intensity).

### Functional assessment of the ELT

Albumin production was measured in supernatant with Human Albumin ELISA Kit (Bethyl Laboratories), according to manufacturer’s instructions. Urea synthesis was measured using the QuantiChrom Urea Assay Kit (Gentaur) according to the manufacturer’s instructions. Briefly, 50 µL of culture medium was transferred into a clear-bottom 96-well plate in duplicate, along with 50 µL of a 5 mg/dL urea standard prepared by diluting the provided 50 mg/dL standard tenfold in culture medium. A blank control consisting of 50 µL of culture medium was also included. Subsequently, 200 µL of freshly prepared working reagent (1:1 mixture of Reagent A and Reagent B) was added to each well and mixed by gentle tapping. The plate was incubated at room temperature for 50 minutes, after which absorbance was measured at 520 nm using a microplate reader. Urea concentrations were calculated using the manufacturer’s equation:

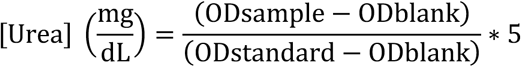

Ammonia was measured in supernatant with Ammonia Assay Kit (Abcam), according to manufacturer’s instructions. Alpha-fetoprotein production was assessed using Alpha-fetoprotein ELISA kit from Abcam according to manufacturer’s instructions. Hepatocytes and HLC were analyzed for cytochrome P450 activity after 8 days of culture and compared to the ELT 4 weeks after encapsulation. The substrate medium was high glucose DMEM without phenol red (Gibco) supplemented with 100 units/mL penicillin (Gibco) and 100 μg/mL streptomycin (Gibco). Substrates were added into medium at proper dilution ratios: CYP3A4-IPA for CYP3A4 (dilution: 1:1000, Promega); 6′deoxyluciferin (Luciferin H) (dilution: 1:50, Promega) for CYP2C9, and Coumarin (50 μm, Sigma) for CYP2A6. The spent medium was removed, and cell aggregates were washed with PBS for 3 times. The cell aggregates were incubated with substrates for 2 hours at 37 °C. After 2 hours, the medium was placed in Eppendorf tubes and frozen at −20 °C for further analysis. Metabolite conjugates formed from coumarin were incubated with β-glucuronidase/arylsulfatase (Roche) for 2 hours at 37 °C. Samples were diluted 1:1 in quenching solution and formation of metabolites was quantified with a fluorescence microplate reader (Molecular Devices) as described elsewhere.^67^ Metabolite conjugates formed from Luciferin H and CYP3A4-IPA were processed and analyzed per Promega protocol and analyzed using a microplate luminometer (Molecular Devices). CYP450 induction or inhibition experiments were performed using rifampin (25 μM), Methoxsalen (3.125 μM), and Sulfaphenazole (25 μM) dissolved in culture medium, and cells exposed for 2 days (Methoxsalen, Sulfaphenazole) or 4 days (rifampin) prior to CYP450 analysis.

### Histological analysis of animal livers

After euthanasia with isoflurane, livers have been quickly harvested from the abdominal cavity, washed in cold PBS, fixed in 4% paraformaldehyde for 12 hours at 4°C, and then left in 70% EtOH until processing. The organs were then cut in half and embedded in paraffin, 10 µm sections were obtained at the microtome, and H&E staining were performed according to standard methods.

### Statistical analysis

All values are shown as mean ± standard error. Mann-Whitney U test paired or unpaired t test with Welch’s correction, one-way or two-way ANOVA and log rank test were used where appropriate. A p-value of <0.05 (two-tailed) was considered significant.

## Abbreviations

ALF: acute liver failure;
DC: dendritic cells;
ELT: encapsulated liver tissue;
EPC: endothelial progenitor cells;
HE: hepatic encephalopathy;
HLC: hepatocyte-like cells;
IgG: immunoglobulin G;
iPSC: induced pluripotent stem cells;
MPC: mesenchymal progenitor cells;
PEG-VS: polyethylene glycol-vinyl sulfone;
PHH: primary human hepatocytes;

## Data availability

The authors declare that the data supporting the findings of this study are available within the paper and its supplementary information files.

## ACKNOWLEDGEMENTS

We thank the *Centre Québécois de génomique clinique* (CQGC, Integrated Centre for Pediatric Clinical Genomics) for sequencing services. We are grateful to Drs. Ariella Shikanov, James Day, and Anu David from the University of Michigan for their advice on PEG–VS encapsulation. We also thank Ms. Ines Boufaied from the CHU Sainte-Justine Flow Cytometry Core Facility for her collaboration.

## FUNDING

This work was supported by the Stem Cell Network grant FY17/DT8 (M.P.), the Canadian Institute of Health Research New Investigator in Maternal, Reproductive, Child & Youth Health grant MY6-155373 (M.P.) and the “Fonds de Recherche du Québec – Santé” Junior 1/2 Clinician-Scientist scholarship (M.P.).

## AUTHOR CONTRIBUTIONS

- Claudia Raggi: conception and design, data collection, analysis and interpretation, manuscript writing.
- Pascal Lapierre: data analysis and interpretation, manuscript writing.
- Marie-Agnès M’Callum: data collection.
- Quang Toan Pham: data collection.
- Silvia Selleri: data collection and analysis and interpretation.
- Radu Alexandru Paun: data collection, analysis and interpretation.
- Nissan Baratang: data collection.
- Chenicka Lyn Mangahas: data collection.
- Basma Benabdallah: data collection, analysis and interpretation.
- Gaël Moquin-Beaudry: data collection, analysis and interpretation.
- Jiwoon Park: data collection.
- Dorothée Dal Soglio: data analysis and interpretation.
- Yves Théoret: data collection, analysis and interpretation.
- Robert E. Schwartz: data analysis and interpretation.
- Christian Beauséjour: conception and design, data analysis and interpretation.
- Elie Haddad: conception and design, data analysis and interpretation.
- Massimiliano Paganelli: conception and design, data analysis and interpretation, manuscript writing, financial support, overall supervision.

All authors approved the final manuscript.

## COMPETING INTERESTS

C.R. and M.P. are co-inventor in patent applications protecting the intellectual property described here, as well as co-founders, shareholders and board members of Morphocell Technologies Inc. P.L and N.B. are employees of Morphocell Technologies. R.E.S. is on the scientific advisory board of Miromatrix Inc. and Lime Therapeutics and is a paid consultant and speaker for Alnylam Inc. and Novo Nordisk.

## Supplementary figures

**Supplementary Figure 1.**
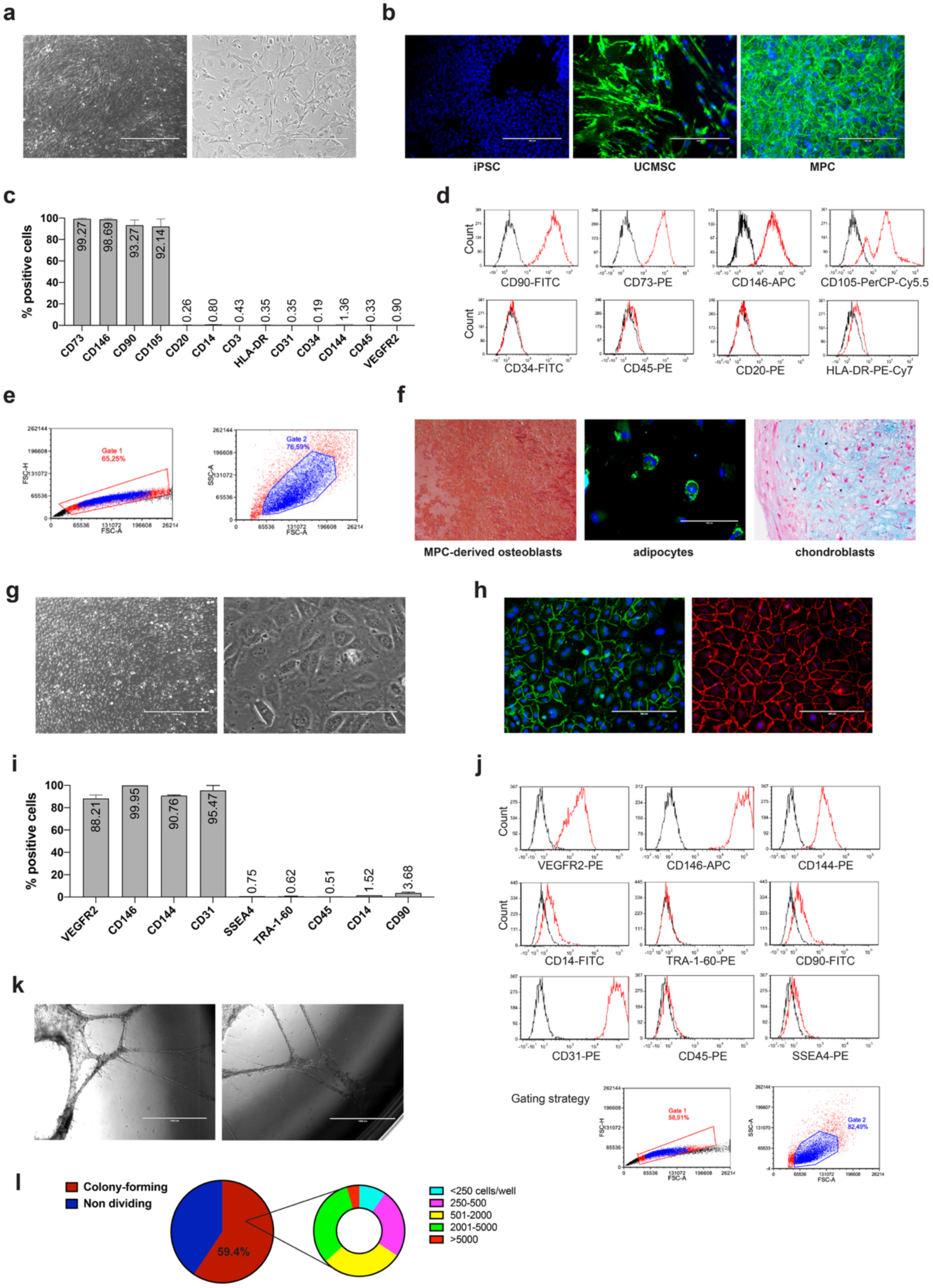
iPSC-derived mesenchymal progenitor cells (MPC) and endothelial progenitor cells (EPC). (Adapted from Raggi et al. Stem Cell Reports. 2022**;17:584-598.) (a)** MPC showed adherence to plastic when maintained in standard culture conditions, with a microscopy appearance undistinguishable from other mesenchymal stromal cells (MSC; phase contrast, representative images, scale bar 1000 µm left, 200 µm right). **(b)** Fibronectin expression (green) by MPC compared to iPSC and umbilical cord-derived MSC (UCMSC; immunofluorescence, representative images of 3 biological replicates, DAPI nuclear staining, scale bar 200 µm). **(c-d)** MPC express surface antigens characterizing MSC (CD73, CD105, CD90) and are negative for markers of other cell types (leukocytes, lymphocytes, antigen-presenting cells, hematopoietic progenitors, endothelial cells) most likely to contaminate MSC cultures (flow cytometry; c: 3 independent differentiations; d: representative graphs). **(e)** Gating strategy used for flow cytometry (representative image of 3 independent experiments). **(f)** MPC could be successfully differentiated into osteoblasts (left, Alizarin red staining), adipocytes (center, LipidTox lipid staining, scale bar 200 µm) and chondroblasts (right, Alcian blue staining; 3 differentiations for each line, representative images). **(g)** Representative morphology of EPC (3 biological replicates, phase contrast, scale bar 1000 µM left, 100 µM right); **(h)** CD31 staining (green and red) in EPC (right) compared to human umbilical vein endothelial cells (HUVEC, left; DAPI nuclear staining; scale bar 200 µM). **(i-j)** Characterization based on positive (CD31, VEGFR2, CD146, CD144) and negative (SSEA4, CD45, CD14, TRA-1-60, CD90) surface markers (flow cytometry, c: summary data from 3 independent differentiations; d: representative graphs). The gating strategy used is shown at the bottom (representative image of 3 independent experiments). **(k)** iPSC-derived EPC spontaneously form tubes when seeded on Matrigel (tube formation assay, representative phase contrast images, scale bar 1000 µM). **(l)** EPC are capable to effectively form colonies after single cell sorting (colony forming assay, representative experiment, lower count threshold 250 cells).

**Supplementary Figure 2.**
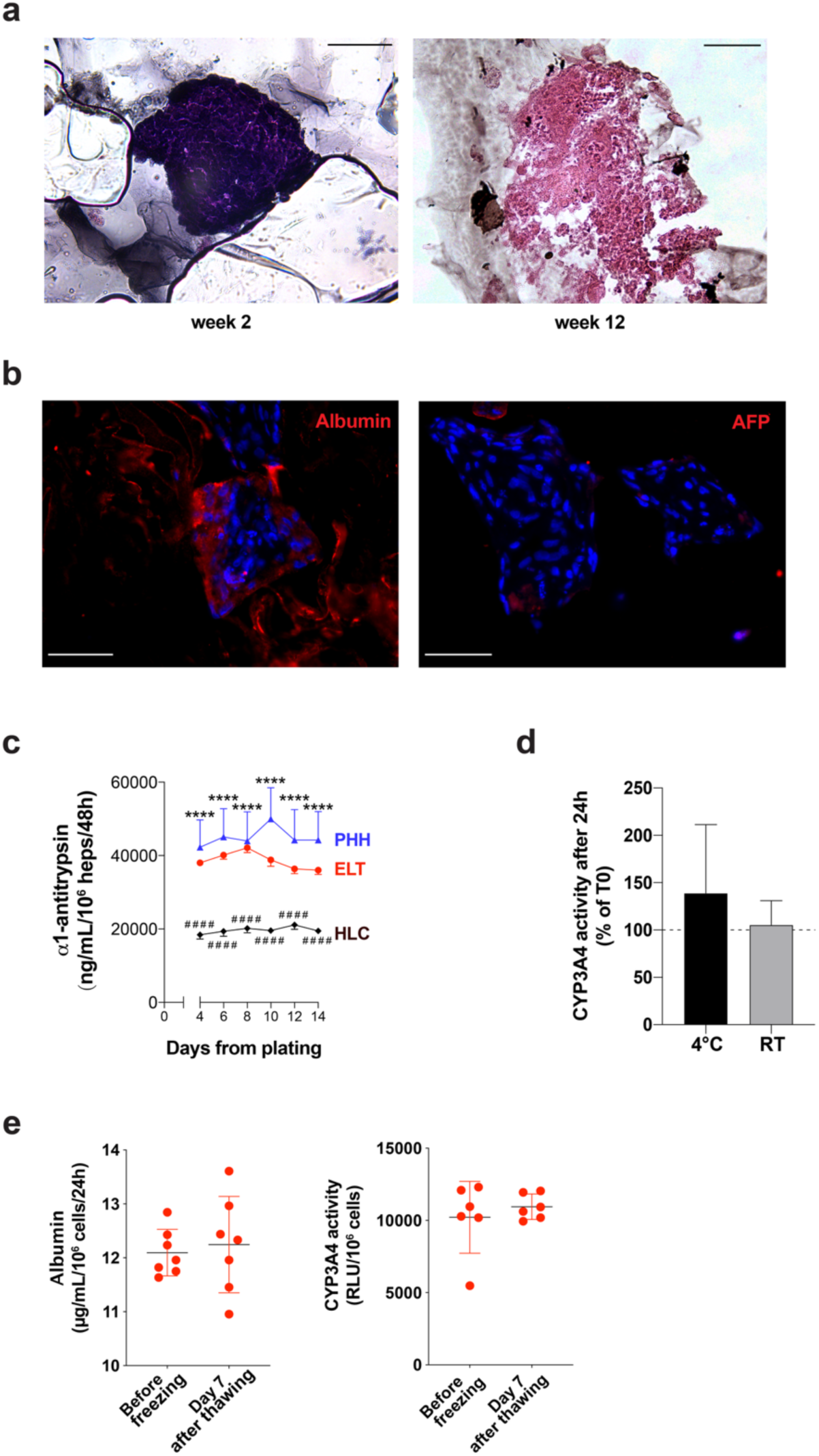
*In vitro* characterization of the ELT. **(a)** Although encapsulation into PEG-VS significantly complicates processing of the ELT samples for histological analysis, no significant growth or change in the morphology of the organoids was observed over time (H&E, representative images, scale bar 50 µm). **(b)** After 2 weeks from encapsulation, the ELT showed strong albumin expression but an already very mild AFP expression (immunofluorescence, representative images, DAPI nuclear staining, scale bar 50 µm). **(c)** The ELT resulted superior to HLC and more similar to PHH in its capacity to secrete alpha1-antitrypsin (ELT n=38 from 6 batches, HLC n=10, PHH n=12; Mann-Whitney test, **** *p*<0.0001 vs PHH, #### *p*<0.0001 vs HLC). **(d)** The ELT was not significantly affected by temperature changes, with liver functions not significantly impaired after 24h at 4°C (shipping test) or even room temperature (Cyp3A4 activity expressed as percentage change compared to standard culture conditions [T0]; n=4, paired *t* test, p=ns). **(e)** The ELT tolerated cryopreservation (−80°C) without significant effects on liver functions (albumin secretion, left and Cyp3A4 activity, right, before and after a freeze/thaw cycle; 1 week at −80°C, n=7 biological replicates, paired *t* test, p=ns).

**Supplementary Figure 3.**
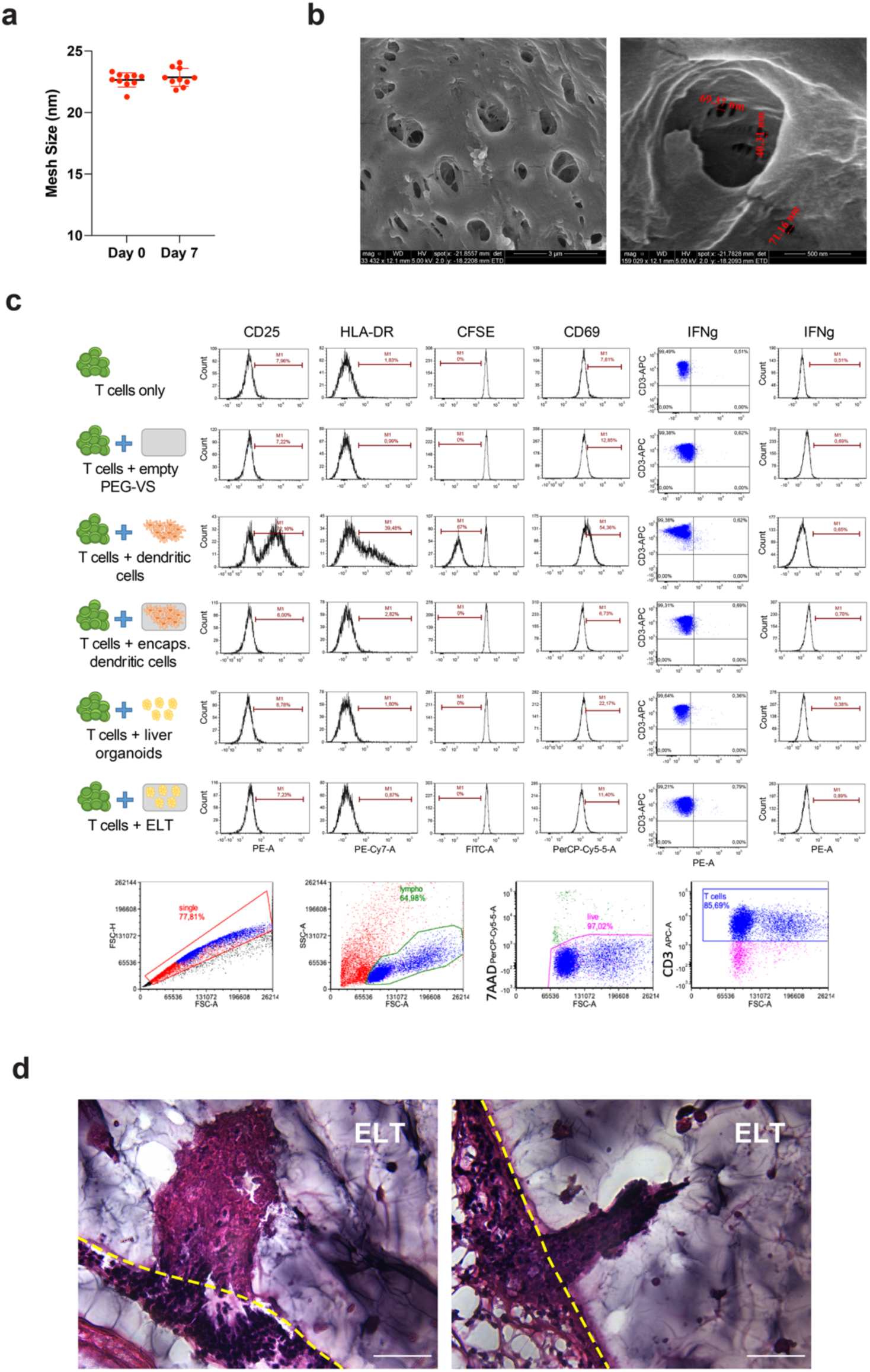
Encapsulation of human ELT protects against rejection. **(a)** The mesh size of 4% PEG-VS did not change upon a freeze/thaw cycle (1 week at −80°C in University of Wisconsin’s solution; n=10, *p*=ns). **(b)** The pore size of 4% PEG-VS was larger at the surface of the biomaterial (left) and decreased in size within the hydrogel to approximately 20 nm (scanning electron microscopy, scale bar: left 3 µm, right 500 nm). **(c)** MLR showed that encapsulation within 4% PEG-VS prevented T cell activation (experimental set up and % of T cells at flow cytometry, representative graphs of the results showed in Fig. 3d; DC: activated dendritic cells, CFSE: carboxyfluorescein succinimidyl ester; n=5 biological replicates with peripheral blood mononuclear cells from 5 different donors). An example of the gating strategy used is shown at the bottom (representative image). **(d)** Without double encapsulation, since encapsulation is obtained by resuspending the organoids within the liquid hydrogel before photopolymerization, a few organoids remain exposed on the surface of the ELT. Such organoids are attacked by the recipient immune system, but the immune cells cannot spread further into the biomaterial (single-encapsulated human ELT explanted after 30 days into the peritoneal cavity of immunocompetent mice; H&E, representative images of 4 biological replicates, scale bar 100 µm).

**Supplementary Figure 4.**
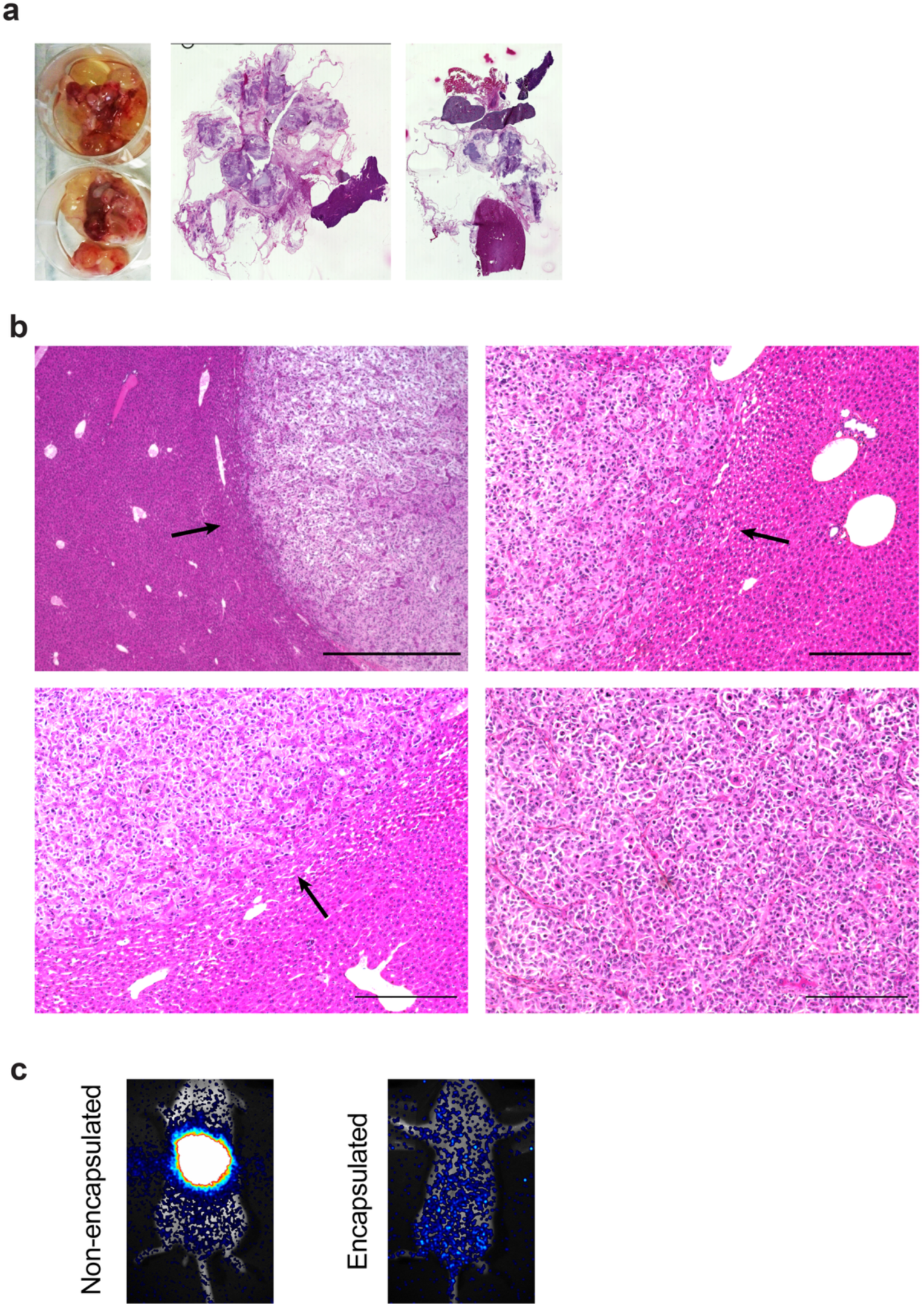
Encapsulation of human ELT protects against tumor formation. **(a)** When implanted into immunodeficient NSG mice, without encapsulation, undifferentiated iPSC generated teratomas after 6 weeks (macroscopic, left and microscopic, left and right, appearance, representative images of 4 biological replicates). **(b)** When injected into the liver of NSG mice, transformed iPSC-derived hepatoblasts gave rise to undifferentiated hepatoblastomas (arrows; H&E, representative images of 6 biological replicates, scale bar 1000 µm left and 300 µm right).**(c)** Encapsulation protected against the development of malignant liver cancer: no tumor formed in NSG mice after 16 weeks when GFP/luciferase-tagged hepatoblastoma cells were encapsulated into 4% PEG-VS before implantation, as compared to non-encapsulated cells forming tumors upon intrahepatic injection in controls (live imaging, luminescence reading adjusted to maximize sensitivity, representative images).

**Supplementary Figure 5.**
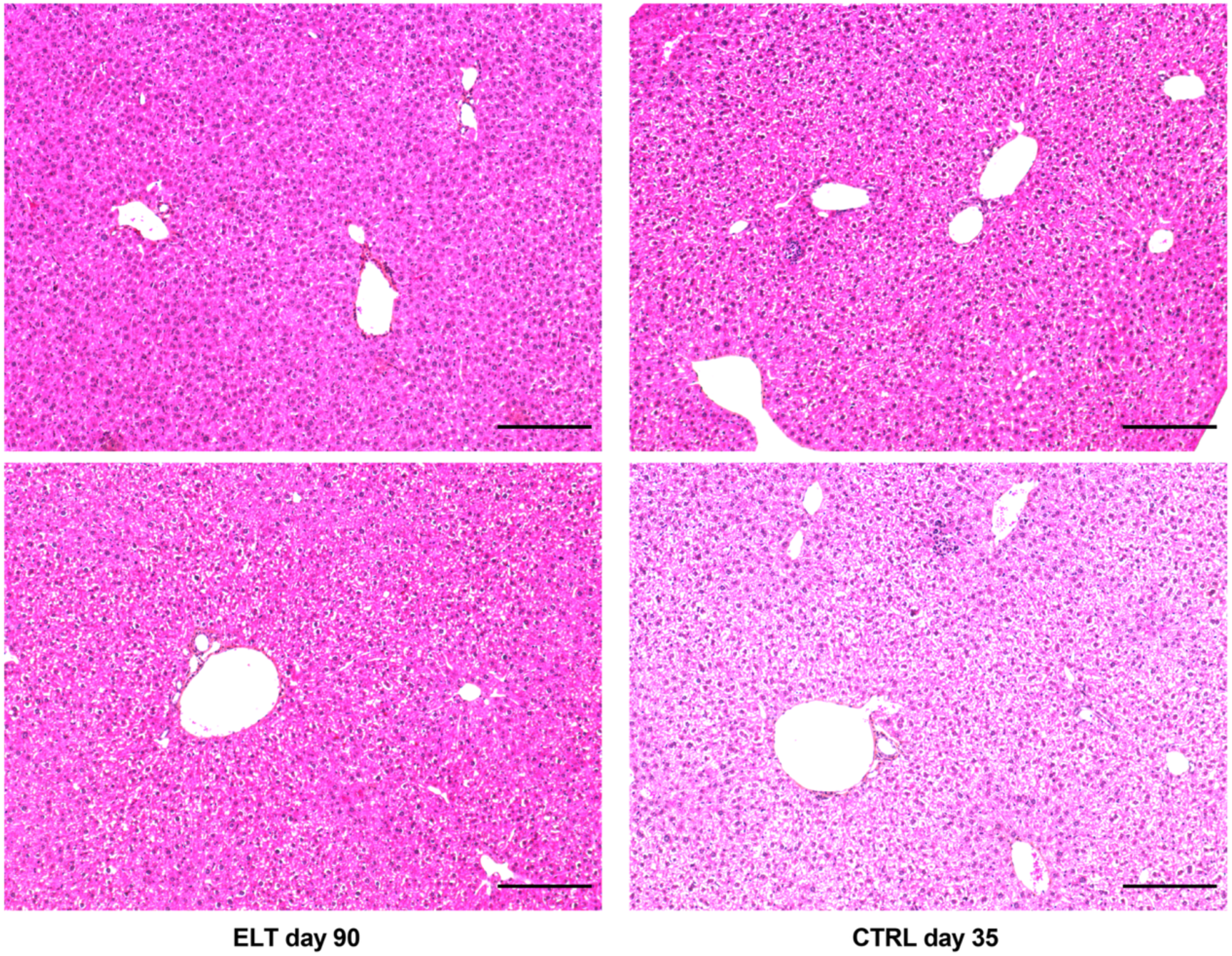
Long-term follow-up of treated mice. Mice with CCl4-induced ALF were sacrificed at day 35 (CTRL, n=5) and day 90 (ELT, 60 days ELT explant, n=12 animals). Livers of survivors from both groups were normal at histological examination (H&E, representative images, scale bar 200 µm).

